# Asymmetric effects of environmental filtering on the assembly of tropical bird communities along a moisture gradient

**DOI:** 10.1101/251249

**Authors:** Juan Pablo Gomez, Scott Robinson, Jose Miguel Ponciano

## Abstract

The species-sorting hypothesis (SSH) states that environmental factors influence local community assembly of metacommunities by selecting for species that are well adapted to the specific conditions of each site. Along environmental gradients, the strength of selection against individuals that are marginally adapted to the local conditions increases towards the extremes of the environment where the climate becomes harsh. In rainfall gradients, the strength of selection by the environment has been proposed to decrease with rainfall. Under this scenario SSH would predict that immigration of individuals from the metacommunity should be restricted into the dry end of the gradient creating a positive relationship between immigration and rainfall. However, if the selection is strong in both ends of the gradient, the restriction should be expected to be in both directions such that the ends behave as independent metacommunities even in the absence of geographical barriers. In this study we used models based on neutral theory to evaluate if SSH can explain the distribution of bird species along a steep rainfall gradient in Colombia. We found a strong positive relationship between immigration rates and precipitation suggesting that the dry forests impose stronger challenges for marginally adapted bird populations. However, a two-metacommunity model separating dry and wet forests was a better fit to the observed data, suggesting that both extremes impose strong selection against immigrants. The switch from the dry forest to the wet forest metacommunities occurred abruptly over a short geographic distance in the absence of any apparent geographic barrier; this apparent threshold occurs where the forest becomes mostly evergreen. The relative number of rare species in dry forest was lower than in wet forests suggesting that the selection against marginally adapted populations is stronger in the dry forests. Overall, our analyses are consistent with SSH at the regional scales, but the rarity analysis suggests that the mechanisms at the local scales are substantially different. Based on these results, we hypothesize that abiotic (climatic) factors limit immigration into dry forest communities and whereas biotic factors such as competition and predation may limit immigration into bird communities in the wet forest.

## Introduction

Environmental factors have long been hypothesized to determine species composition of a community by filtering out species that are not adapted to the particular conditions of the locality (Grinell 1917, Weiher and Keddy 1999, Condit et al. 2002, Tuomisto et al. 2003). If species abundances are used as surrogates of their fitness under local environmental conditions (Sokol et al. 2011), then the abundances of species should change as environmental conditions change through climatic space (Chase and Myers 2011). Viewed in this context, environmental gradients can be treated as natural experiments in which the role of environmental conditions in community assembly processes can be tested under varying conditions (McGill et al. 2007, Chase and Myers 2011).

Along environmental gradients, the spatial scale can be small enough that no geographical barriers limit dispersal between local communities. In these types of environmental gradients, local communities are connected through dispersal and immigration, homogenizing the local communities at broad spatial scales. This group of local communities can be treated as a metacommunity (sensu Leibold et al. 2004), which is defined as a set of local communities that are connected through dispersal. Within a metacommunity, the assembly process is the result of two independent steps (Etienne 2007, Munoz et al. 2007, Jabot et al. 2008, Etienne 2009): 1) formation of species in the metacommunity and 2) immigration/recruitment in the local community (Jabot et al. 2008). Within local communities, the assembly process reflects a zero-sum game, in which an immigrant from the metacommunity or a recruit from a species already present in the local community immediately replaces every individual that dies (Hubbell 2001).

In metacommunities distributed among homogeneous climates, the assembly process could be mainly determined by dispersal limitation and biotic factors (e.g. competition, parasitism). Limitations to dispersal should be a function of the distance of a particular community to the rest of the sites (Hubbell 2001). However, in metacommunities distributed along environmental gradients, the restriction to dispersal should be additionally influenced by the abilities of the immigrants to recruit in the climatic conditions of each site (Weiher and Keddy 1999). All individuals might be able to disperse into the local communities, but only individuals from species that are adapted to the local climatic conditions can recruit successfully (Grinell 1917, Weiher and Keddy 1999, Jabot et al. 2008). Those species that fail to recruit in a local community may appear as if they were unable to disperse, even though their absence actually reflects their inability to become established. This process has been referred to as the species-sorting hypothesis (SSH; Leibold et al. 2004).

Along environmental gradients, the selection against immigrants is likely to be stronger at the extremes of the gradient where climates are likely to be harsher. In the extremes, SSH predicts that, each time an individual is lost from the community it is more likely to be replaced by an individual of a species that is already present in the local community than it is by an immigrant from the metacommunity. Harsh conditions therefore should decrease immigration from the metacommunity (Leibold et al. 2004). In addition, it is likely that species colonize localities with one or few individuals. If such species are marginally adapted to extreme environmental conditions, strong selection by the environment would likely eliminate entire populations of these species, favoring the recruitment of already established, well-adapted and abundant species. Communities in harsh environments, therefore, should have reduced species diversity (few rare species and more common species), and should show greater differentiation from the greater metacommunity regardless of the geographic distances separating these communities from those in less harsh environments. This latter prediction is identical to the prediction of dispersal limitation by distance except that in this case, differentiation should be proportional to the differences in climatic variables rather than geographic distance *per se*.

Rainfall gradients are hypothesized to have stronger selection at the drier end, possibly because of limited water availability and greater temperature variability (Condit et al. 2002, Engelbrecht et al. 2007, Jabot et al. 2008, Graham et al. 2009). In this case, SSH would predict that immigration of individuals from the metacommunity to localities in the dry forest should be reduced as a result of the inability of most wet forest species to recruit in dry forests. In the wet end, where climatic conditions are more benign, the immigration of an individual should not depend on its ability to overcome the environmental conditions, which should lead to higher immigration rates from the metacommunity and greater persistence of species with small populations. Thus, the immigration rate should increase with precipitation as individuals have fewer restrictions on immigration into the wetter end of the gradient (Jabot et al. 2008). Alternatively, if there is no effect of climate on the immigration process, then local communities should have similar immigration rates or at least there should not be a relationship between precipitation and immigration rates (Jabot et al. 2008). Instead, immigration rates should be lower in local communities that are at the periphery of the metacommunity (Hubbell 2001). Empirical values of immigration to local communities are difficult to obtain even at small spatial scales. However, several models based on Neutral theory allow the estimation of immigration parameters from observed abundance data collected in the field (Etienne 2007, Munoz et al. 2007, Jabot et al. 2008, Etienne 2009). Even though these models are essentially neutral, the reinterpretation of the immigration parameter into a recruitment limitation parameter allows for testing the role of environmental filtering in community assembly (Jabot et al. 2008).

Even though bird communities were very important in the development of current community ecology theory (Preston 1948, Cody 1974, Diamond and Cody 1975), plant systems have been mostly used to test the effect of environmental conditions on community assembly and the development of neutral models (Weiher and Keddy 1999, Hubbell 2001, Condit et al. 2002, Tuomisto et al. 2003, Engelbrecht et al. 2007). Birds, however, might be a useful system to test the effects of climate on species abundance distributions, because their dispersal abilities might be greater and because they can actively select their habitat and environments. Thus, neutral processes might be less important for birds than they are for plants, allowing us to increase the power to detect the influence of climatic conditions in community assembly.

The Magdalena River Valley is an inter-Andean lowland valley located between the central and Eastern cordilleras in Colombia, and contains a precipitation gradient that ranges from 700 mm in the south up to 4000 mm in the north, but has little variation in elevation and temperature. There are no geographic barriers that prevent dispersal of species allowing us to make the assumption that the localities within the valley are potentially part of one metacommunity. In this study, we obtained information on the community composition from 13 localities along this gradient; to test if SSH is a good explanation for the observed distribution patterns of birds in the Magdalena Valley in Colombia. Specifically, we test the prediction that individual immigration rates are lower in localities with harsher climate (i.e. dry forests). Alternatively, we ask if the strength of selection is high in both ends of the gradient. In this case, SSH would predict that the localities within the gradient behave as two independent metacommunities (i.e. dry forest and wet forest) even in the absence of geographical barriers separating them. Finally, we test if dry forests have fewer rare species than wet forests, as predicted by the SSH based on reduced immigration rates under harsher environmental conditions.

## Methods

### Bird sampling

We selected 13 localities along the valley to determine community composition. In each of these 13 localities we placed a minimum of nine 50-m fixed-radius point counts (Hutto et al. 1986) in which we censed all birds (Table 1). We located point counts only in forest habitats separated by 200 m and at least 75 m from the edge (Blake and Loiselle 2001). Points were visited from dawn to no later than 10:00 AM avoiding days that were too windy and rainy between the months of June and August 2012 and January and August 2013. At each point count we counted all of the individuals seen and heard within ten minutes.

**Table 1.** Characteristics of the study sites. Classification into the Dry forest or Wet forest ecoregions follows Olson *et al* (2001)

| Locality | Ecoregion | Points | Precipitation | Species | Individuals | # Rare Species |
| --- | --- | --- | --- | --- | --- | --- |
| Venadillo | Dry Forest | 20 | 1224 | 87 | 1483 | 17 |
| Bateas | Dry Forest | 36 | 1280 | 102 | 1868 | 20 |
| ManaDulce | Dry Forest | 32 | 1375 | 117 | 2959 | 14 |
| Potosi | Dry Forest | 30 | 1379 | 110 | 2926 | 15 |
| Jabiru | Dry Forest | 26 | 1618 | 105 | 2314 | 12 |
| Boqueron | Dry Forest | 13 | 1668 | 101 | 2022 | 15 |
| Rio Manso | Wet Forest | 25 | 2240 | 118 | 1980 | 28 |
| Rio Claro | Wet Forest | 14 | 2428 | 141 | 2350 | 38 |
| Maceo | Wet Forest | 13 | 2528 | 133 | 2222 | 25 |
| La Perla | Wet Forest | 9 | 2755 | 121 | 1812 | 29 |
| Remedios | Wet Forest | 20 | 2824 | 126 | 1736 | 28 |
| Barbacoas | Wet Forest | 34 | 2888 | 120 | 1796 | 29 |
| San Juan | Wet Forest | 10 | 2997 | 97 | 1672 | 26 |

### Abundance estimation

We used a zero-inflated Poisson model to estimate the mean number of individuals per species per point count and extrapolated it to 100 ha. Variation in the mean number of individuals among point counts could be modeled using a zero-inflated negative binomial model; however, this model tends to overestimate abundance (Joseph et al. 2009). We selected 100 ha because it allowed comparisons with other studies from tropical lowland forests (e.g. Terborgh et al. 1990, Thiollay 1994, Robinson et al. 2000, Blake and Loiselle 2009). We excluded hummingbirds, parrots, swifts and swallows, nocturnal birds and birds that flew over the point count during the sampling time. All of the estimates were derived using the pscl package (Achim et al. 2008) in R v 3.0.2 (R Core Team 2013). We alternatively estimated the abundance of each species using the N-mixture models with Precipitation as an abundance covariate and time of day and Julian day of the year as a detection covariate (Royle 2004, Joseph et al. 2009). We finally inspected the estimates visually to determine the accuracy of the models to estimate the abundance of each species, and compared them to previous estimates and the knowledge of an expert (Terborgh et al. 1990; S.K. Robinson).

### Model fitting

We used the multiple samples estimation approach (Etienne 2007, Munoz et al. 2007, Jabot et al. 2008, Etienne 2009) to estimate the immigration parameters that best described the data. In this approach, the assembly process is assumed to be a two-step process in which speciation occurs at the metacommunity level, and then local communities are assembled through immigration (Munoz et al. 2007). Within the local communities, the dynamics are assumed to follow a zero-sum game, in which any deaths in the local communities have to be immediately replaced by; (1) a recruit from the local community or (2) an immigrant from the metacommunity. The zero-sum game makes the assumption that all niches in the local community are occupied and that the number of individuals are kept constant. The two-step assembly process is described by two parameters that allow the prediction of the species abundance distribution of each local community. The Fundamental Biodiversity number (θ)(Hubbell 2001), describes the diversity of the metacommunity based on its size (i.e. number of individuals) and the rate of introduction of new species (i.e. speciation rate). The immigration rate (*m*) parameter is defined as the probability that an immigrant from the metacommunity replaces a death in the local community. Low values of *m* indicate high restriction to dispersal because it is more likely that recruits from the local community replace dead individuals. Alternatively, *m* can be reinterpreted as the potential number of immigrants from the metacommunity (I) that compete with local individuals for available sites (Etienne and Olff 2004). *I* and *m* are related as follows: 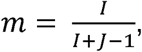 where *J* is the number individuals in the local community.

In the Magdalena Valley, there are no geographic barriers that prevent dispersal of species allowing us to make the assumption that all of our communities are potentially part of one same metacommunity. In the two-step assembly process two scenarios can arise. The first occurs when immigration is constant from the metacommunities to each local community. In this case, a two-parameter model is sufficient to predict all of the species abundance distributions of the 13 localities (i.e. Constant Immigration model; Etienne 2007). The second case occurs when the migration rates are different for each locality in the metacommunity (i.e. Variable Immigration model;Etienne 2009). If the environment is not important for determining the species’ abundances along the valley, then the Constant Immigration model with only two parameters will be a better fit to the data. Alternatively, if the environment is important for structuring communities and the dry forests present a stronger environmental filter, then the Variable Immigration model will fit the data better and the immigration rate should increase with precipitation. Thus, we fitted the models with Constant and Variable immigration using the numerical optimizations presented by Etienne (2007) and Etienne (2009), respectively.

Alternatively, it is possible that selection is strong and both ends of the gradient. This would cause immigration between sites in different habitats to be low, such that the gradient can be split into two or more metacommunities (Leibold et al. 2004). One way to test for this effect is to assume that dry forests and wet forests each represent a separate metacommunity. We used the definition of the WWF ecoregions to group the localities into dry forest and wet forest communities separately (Olson et al. 2001) and fitted the Constant and Variable immigration models to the dry forest and wet forest localities separately. Finally, it is also possible that isolation from the metacommunity results in reduced immigration rates (Hubbell 2001). To account for the effect of distance, we estimated the centroid of the Magdalena Valley metacommunity and calculated the distance from each local community to the centroid. We then used this distance as a measurement of isolation, assuming that local communities that are furthest from the centroid are in the periphery of the metacommuntity and are therefore more likely to receive a lower number of immigrants.

To test for the effects of precipitation and distance on immigration rates (*m*, *I*), we fitted a linear model that had precipitation and distance as predictor variables. We compared the AIC of this model with the models including only precipitation and only distance to determine which variable had the greatest influence on the variation of immigration rates. We then repeated the analysis with the immigration estimates from the variable immigration two-metacommunity model. In the latter case, we calculated the centroid for each metacommunity separately, and found the distance from each local community to the centroid of the metacommunity in which it belongs. We also compared the immigration estimates between the dry and wet forests and between variable immigration one and two metacommunty models using ANOVAs.

We compared the fit of the models using AIC in which the model with the lowest AIC was taken as the best, and models were significantly different if the difference in AIC was greater than 2. To assess the goodness-of-fit of the best model for predicting the species abundance distribution, we fitted a fully parameterized model in which for each local community we estimated a θ parameter as well as an immigration parameter using the Etienne’s sampling formula (i.e. Full model; Etienne 2005). The goodness-of-fit is given by a likelihood ratio test between the full model and the best model (Strong et al. 1999). All of the maximum likelihood estimates were performed following Etienne’s (2007, 2009) modification of the Etienne’s sampling formula (Etienne 2005) and were performed in PARI/GP v. 2.7.0 (The PARI Group 2014) using the programs available in the supplementary information of Etienne (2007, 2009).

The 95% confidence intervals for the parameters of the models were estimated using parametric bootstrap. Because of computational limitations, we only estimated confidence intervals for the best two models. We simulated 1000 communities using the maximum likelihood estimates of the observed data and the observed size of the metacommunity and each of the local communities. The simulations were performed following the urn procedure of the two-step approach suggested by Etienne (2007).

### Rarity analysis

Because habitat filtering should reduce the level of immigration into local communities, each time an individual is lost from the community, it is more likely to be replaced by an individual of a species that is present in the site than by an immigrant from the metacommunity. For this reason, communities that are structured by environmental filtering should have fewer rare species than communities in which other factors are more important during community assembly. To test this, we counted the number of species that had 2 or fewer individuals per 100 hectares in each community and related the number of rare species to precipitation. Because wet forest habitats usually have more species, the increase in the number of rare species can be due to differences in overall number of species per locality. To account for this potential bias, we constructed a model to explain differences in the number of rare species in which we assumed that the sampling error is Poisson distributed, considering precipitation and species richness together and also separately. We then compared the models using AIC. We repeated the analysis setting rarity to: < 5 individuals/100 ha.

## Results

### Abundance estimation

We estimated the abundance of a total of 222 species of birds along the entire valley. The estimates based on a zero-Inflated Poisson model for the abundance of each species resulted in realistic estimates (Table S1). The N-mixture models resulted in an overestimation of the abundance by one order of magnitude (results not shown); thus, we used for our analysis the results of the zero-inflated-Poisson model. Detailed description of the localities is presented in Table 1.

### Comparison between models

The model that fitted the data best was Variable immigration with two-metacommunities, with high goodness-of fit (Table 2; see Table S2 for parameter estimates for all models; Likelihood ratio test: X^2^ = 9.52, df = 12, p =0.3). The second best model was the Variable Immigration with one meta-community (Table 2), but the low goodness-of-fit was low (Likelihood ratio test: X^2^ = 9.52, df = 12, p =0.3). Immigration estimates were significantly higher in the variable immigration two-metacommunity model than the one-metacommunity model (Fig 2; ANOVA: m: F = 53.24, df = 24, p<.001; I: F = 21.27, df = 24, p<0.001).

**Table 2.** Model Selection for the 13 metacommunities in the Magdalena Valley. We present the number of parameters per model, the log likelihood of the Akaike Information Criterion (AIC) and the difference in AIC between the four models fitted to the data. The Full model is the model with different θ and restriction to dispersal (*m*<1), Constant Immigration and Variable Immigration models consider one θ such that the Magdalena Valley would be considered as one metacommunity (One Metacommunity) and two θ such that Dry forests and Wet forests are considered independent metacommunities but with constant and variable immigration rates respectively. Bold indicates the best model by the differences in AIC.

| **Models** | **Parameters** | **LogLik** | **AIC** | **deltaAIC** |
| --- | --- | --- | --- | --- |
| *Full Model* | 26 | -1287.26 | 2620.52 | 6.48 |
| ***Constant Immigration*** |  |  |  |  |
| *One Metacommunity* | 2 | -6166.6 | 12337.2 | 9723.16 |
| *Two Metacommunities* | 4 | -4481.92 | 8971.84 | 6357.8 |
| ***Variable Immigration*** |  |  |  |  |
| *One Metacommunity* | 14 | -1314.38 | 2656.75 | 42.72 |
| *Two Metacommunities* | **15** | **-1292.02** | **2614.04** | **0** |

### Variable Immigration One-metacommunity model

Immigration rates showed a positive significant relationship with precipitation (m: F = 19.57, df = 11, p > 0.001, r^2^ = 0.607, m = 0.007 + 9*10^−3^\**Precipitation*; I: F = 10.75, df = 11, p < 0.01, r^2^ = 0.49, I = 23.6 +14.7\**Precipitation*). Distance of the locality from the metacommunity had little influence on immigration estimates (Table S3). Potential immigrants significantly decreased with distance from the centroid; the best model was the one including both distance and precipitation (F = 10.41, df = 11, p < 0.01, r^2^ = 0.67, m = 0.005 + 0.009\**Precipitation* + 6.9 × 10^−6^\**Distance*; F = 7.3, df = 11, p = 0.01, r^2^ = 0.59, I = 39 + 12\**Precipitation* + 0.04\**Distance*). However, the proportion of variance explained by the Precipitation term was considerably higher than the variance explained by the Distance term and the distance coefficient resulted to be non-significant (m: Precipitation = 67.2, Distance = 0.33; I: Precipitation = 49.4%; Distance = 9.9 %). After correcting for the effects of distance on the immigration estimates, local communities in the dry forests had significantly lower immigration estimates than local communities in the wet forests (Fig 2; ANOVA: m: F = 30.23, df= 11, p < 0.001; I: F = 9.32, df = 11, p = 0.01).

### Variable Immigration Two-metacommunity model

The relationship between immigration and precipitation was not significant (Fig1; *m*: F = 0.98, df = 11, p = 0.34; *m* = 0.048 – 2.6*10-^3^\**Precipitation*; *I*: F = 3.04, df = 11, p = 0.1; I = 121.9 – 14.6\**Precipitation*). Distance had no influence on the immigration estimates for the local communities (Table S3). After correcting for distance, local communities in the dry and wet forests showed similar immigration estimates (Fig 2; ANOVA: m: F = 1.85, df = 11, p=0.2; I: F=4.43, df = 11, p=0.06).

**Figure 1.**
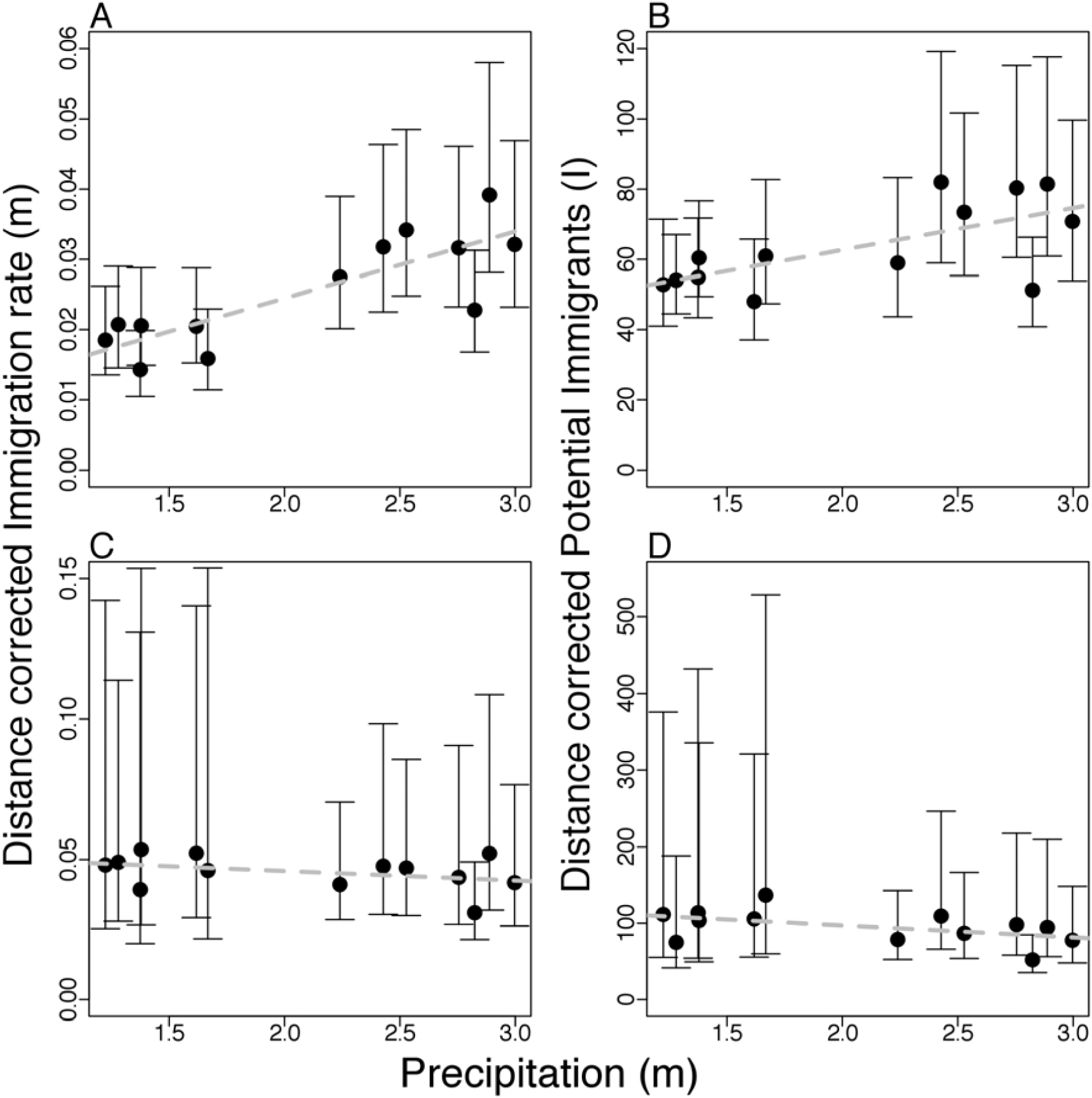
Effects of precipitation on the immigration rates (*m*) (A, C) and potential number of immigrants (*I*) (B, D). The two top panels refer to the one metacommunity model and the two bottom panels show results from the two-metacommunity model. Immigration rates (*m*), are the probabilities that the open space in the local community is occupied by an immigrant from the metacommunity. Potential number of immigrants (I) is the available number of immigrants from the metacommunity that compete for a space open in the local community with recruits of individuals from the local community. The values presented account for the effects of distance from the centroid of the metacommunity in dispersal limitation by using the following correction: *Immigration Estimate* – *β*_2_ *Distance* = *A* + *β*_2_ *Precipitation*, where *β*_1_ and *β*_2_ are the coefficients for Distance and Precipitation, respectively, and *A* is the intercept of the model.

**Figure 2.**
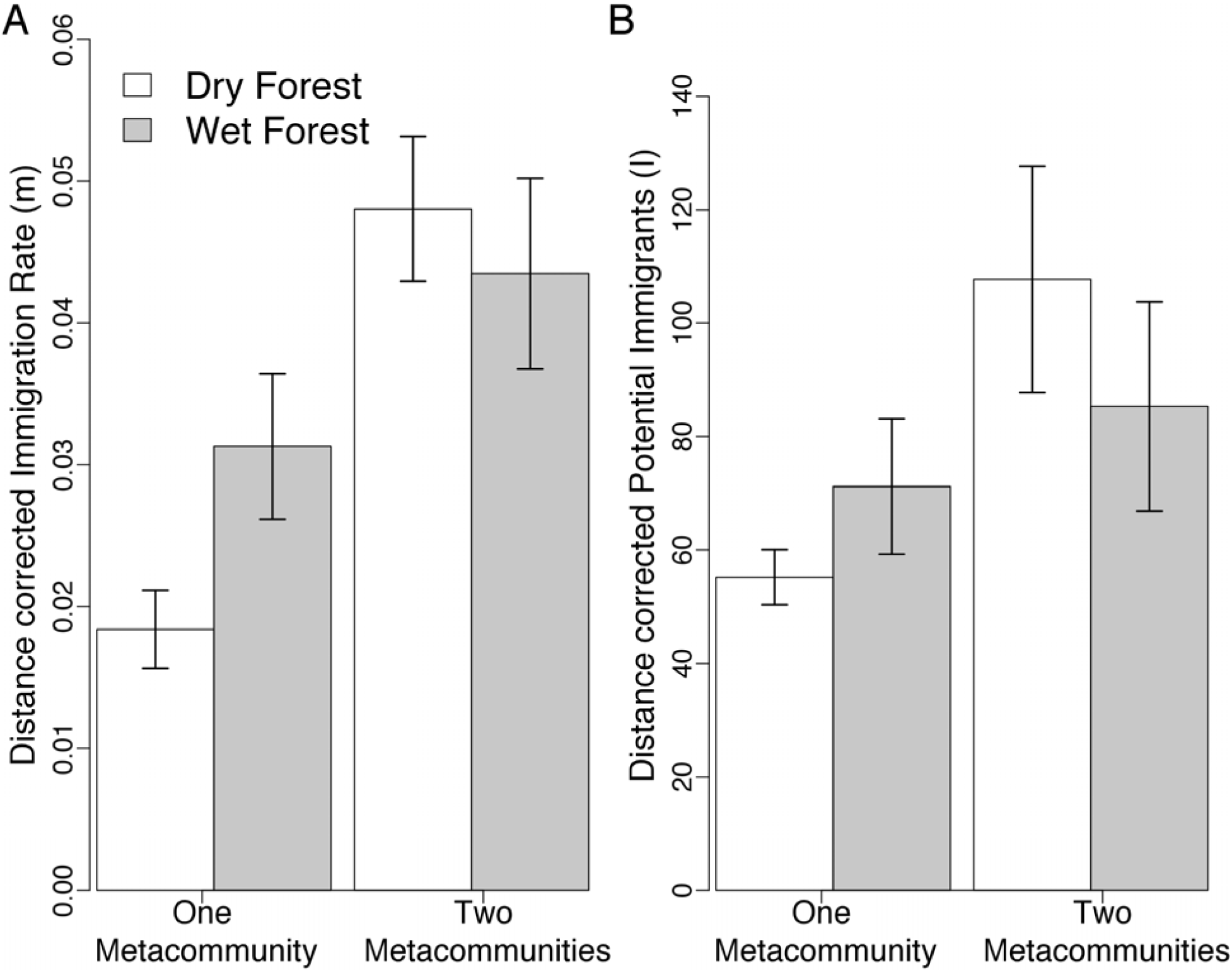
Differences in A) Immigration rates (*m*) and B) Potential number of immigrants (*I*) between types of forest and different models used to estimate immigration. The error bars show the standard deviation of the mean parameter in each type of forest.

### Rarity

In the 2 ind/ha case the best model was the one including precipitation and species richness (*rare species* (2 *ind/ha*) = *e*^(1.6+0.33\**Precipitation*+0.006\**Richness*)^); however, the richness term was non-significant (Table S4). In the 5 ind/ha case the best model was the one including precipitation only (*rare species* (5 *ind/ha*) = *e*^(3.2+0.2\**Precipitation*)^) but was similar to the full model (Table S4). However, in the precipitation plus species richness model, the latter term was not significant (Table S4).

Because species richness had a positive influence on the number of rare species in the analysis with 2 ind/ha, we applied the following correction: 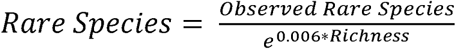. After applying the correction, rarity in the local communities increased from the dry forest to the wet forest (Figure 3). The analysis suggested that the model with different mean numbers of rare species per metacommunity was more likely than the model with no differences between types of forests in both the 2 ind/ha and 5 ind/ha cases (Magdalena = 11/39, logLik = -31.8/-47.1; Dry = 8/32, Wet = 14/46, logLik = -29/-39; X^2^ = 8.7/14.3, p < 0.01 in both cases).

**Figure 3.**
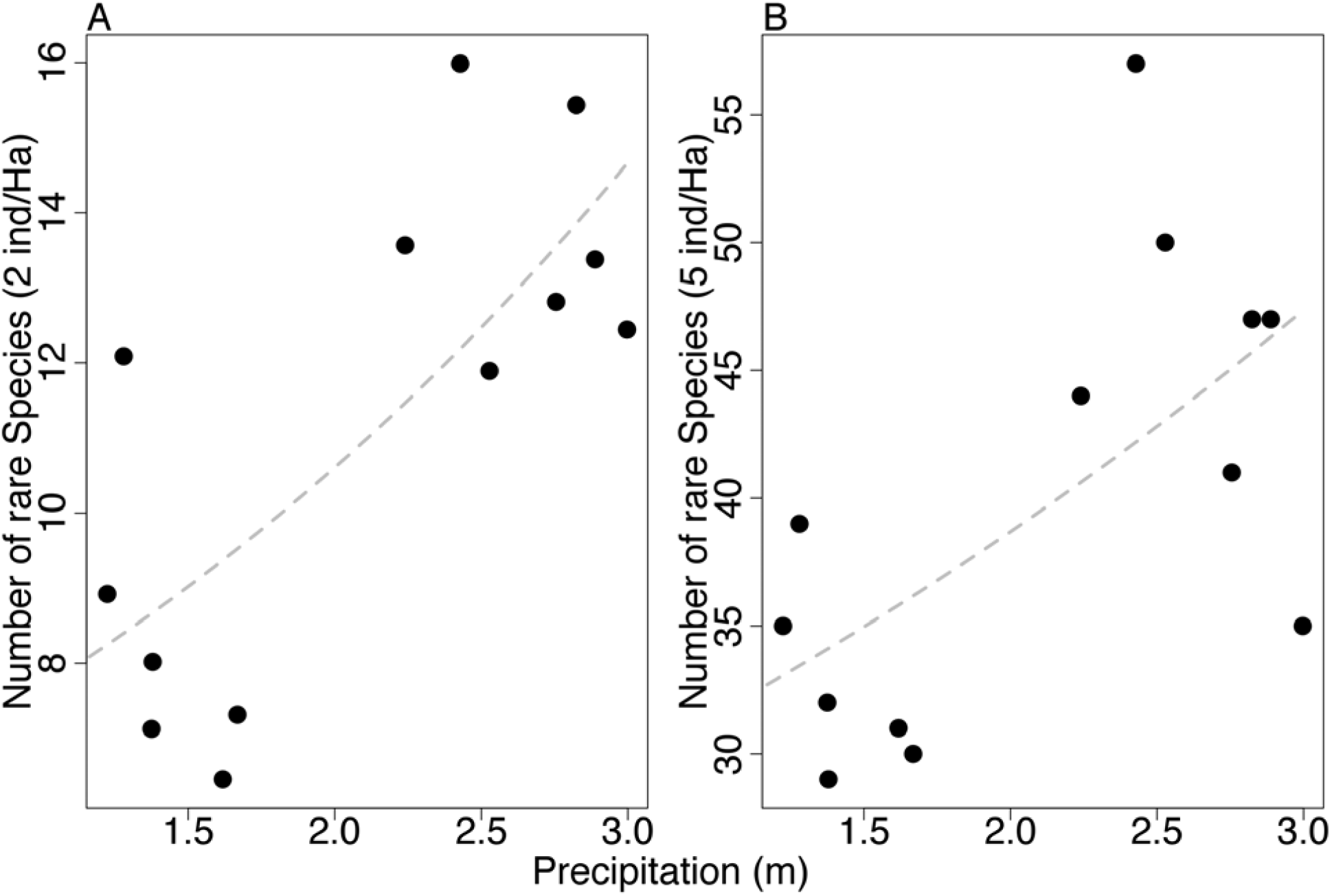
Relationship between precipitation and number of rare species in the Magdalena Valley. A) shows the results for the species with less than 2 ind/ha and B) shows species with less than 5 ind/ha.

## Discussion

Bird communities along the precipitation gradient in the Magdalena Valley can essentially be divided into two metacommunities, even in the absence of geographic barriers. This result suggests that there is strong immigration limitation from dry forest to wet forests and vice versa. However, immigration rates from a metacommunity consisting of both dry and wet forest birds is lower in the dry forest, suggesting that wet forest birds might be less able to immigrate into dry forests than the contrary. This pattern is consistent with the SSH, which predicts reduced immigration of birds into the harsher conditions of the dry forest. There were also indications that dispersal of birds from dry to wet forests was limited, but the effect was smaller (Figure 2). The relationship between immigration rates and precipitation was negative when estimating the immigration rates from dry and wet forest metacommunities independently, which indicates that individuals move more freely among sites in the dry than in the wet forest. Once a species colonizes the dry forest habitat it can invade any locality within the dry forest, but the opposite might not be true in wet forests. Thus, habitat filtering appears to operate mainly at the regional scale through exclusion of wet forest species from dry forest environments, but alternative mechanisms such as biotic constraints (see below) might operate at a local scale in wet forests.

There is little evidence that distance strongly affects similarity among communities, a result suggesting that dispersal limitation has little effect on these communities as suggested by Hubbell (2001). The abundance of bird species remain almost constant across the Magdalena Valley within the same type of forest, a similar result to what Terborgh et al (1996) documented in Amazonian tree communities. Distance from the metacommunity also had little explanatory power for the observed variation in the immigration estimates, which further suggests that dispersal limitations does not affect community organization. Furthermore, the fact that the two-metacommunity model better fits the data suggests that the forest transition zone, which is only about 20 km wide in the Magdalena Valley, drastically affects the abundance of bird species. Thus, there seems to be a threshold effect in which communities shift from a dry forest to a wet forest state over a short distance, rather than a gradual change as precipitation increases.

Rare species seem better able to persist in wet than in dry forests, a result suggesting a relaxation of the selection pressures that would allow immigrants of new species to establish and maintain populations that are rare initially. In contrast, only a subset of species from the metacommunity can immigrate into dry forests, perhaps because the selection pressure is stronger, which would only allow species that can tolerate the harsh climate to establish populations. Under more benign conditions, selection may not be strong enough to eliminate the rare species that are only marginally adapted to the environment, or that have highly specialized niches that could not be maintained in drier forests. Such species could be considered as occasional species that come and go in communities during the dynamic process of community assembly (Magurran and Henderson 2003). However, in dry forests where selection is more intense, marginally adapted, occasional species cannot persist long enough to be captured in the snapshot time frame of typical community studies such as this one.

One potentially strong filter that immigrants would have to overcome in the dry forest is high daily temperature variability. Data logger measurements across two years in a dry and a wet forest locality showed that the coefficient of variation of the dry forest was at least twice that in the wet forest (Coefficient of Variation[CV] in Temperature in dry forest = 0.14, CV in wet forest = 0.006; J.P. Gomez, *unpublished data*). Such extreme variation could impose a challenge to recruitment, especially in early life stages when eggs and nestlings are particularly susceptible to temperature changes (Webb 1987). Wet forest species, however, might be able to recruit in dry forests along creeks and in riparian habitats where climate is much less variable than in the surrounding dry forest. In such habitats, immigrants can overcome the selection by the environment and establish viable populations at low abundances. In fact, most of the wet forest species that that inhabit the dry forest were found along creeks or in riparian vegetation (e.g. *Leptopogon amaurocephalus*, *Mionectes oleagineus*; Table S1).

An alternative interpretation to the patterns of rarity that we documented is that the mechanisms underlying abundance are substantially different in the wet end of the gradient. While dry forests allow only the immigration of well-adapted species to the harsh climate, wet forests allow the immigration of a larger set of species, some of which may depend upon resources and microhabitats that are not available in dry forests. The abundance of such species that specialize on these resources, however, might be further constrained by other mechanisms such as competition (e.g. *Gymnocichla nudiceps*, *Gymnopythis bicolor*; Table S1; Robinson and Terborgh 1995, Touchton and Smith 2011) or mutualisms such as mixed-species flocks (e.g. *Myrmotherula axillaris*, *Epinechrophylla fulviventris*; Greenberg and Gradwohl 1986, Martinez and Gomez 2013). Some of the rare species in the community may also be limited by increased interspecific competition in the more diverse wet forest communities. The abundance of the Spotted Antbird has been shown to be determined by the presence of Ocellated Antbirds, which are behaviorally dominant at ant swarms (Willis 1973). When released from this competition by the extinction of the Ocellated Antbird, the Spotted Antbird has been shown to increase greatly in abundance, suggesting that its rarity was previously maintained by competition (Touchton and Smith 2011). Russo et al (2003) also showed that smaller species are often less abundant than larger species in some foraging guilds in tropical wet forests. The previous result might be attributed to their subordinate status during aggressive interactions that might limit their access to clumped resources (Robinson and Terborgh 1995). It is also possible that the more diverse community of nest predators available in wet forests may further constrain populations of some species (Ricklefs 1969, Martin 1996). Therefore, in the relatively more stable climatic conditions of wet forests, we might predict a greater role of biotic interactions such as mutualisms, nest predation, and competition in structuring communities.

Several other studies have shown effects of environmental filtering on the assembly of tropical communities. The precipitation gradient along the Panama Canal shows evidence of reduced immigration of plant species into the dry forest (Jabot et al. 2008). There is less similarity among plots along this 200-km gradient than there is over thousands of kilometers in Amazonia (Condit et al. 2002). At a global scale, Jabot and Chave (2011) analyzed density-dependent parameters at the community level and concluded that environment factors might govern the assembly of communities. They showed that in regions with low precipitation, the species with higher abundances have a higher probability of recruiting individuals than species with lower abundances. This can be interpreted as a positive density-dependent effect related to the fitness of a species in particular environmental conditions (Sokol et al. 2011).

The effect of environmental filtering on structuring bird communities has mainly been demonstrated in temperate regions, where the climate is believed to be harsher than in tropical environments (Meynard and Quinn 2008, White and Hurlbert 2010, Ozkan et al. 2013). In tropical regions, Graham and coworkers (2009, 2012) demonstrated that dry forest hummingbird communities are composed of species that are closely related, suggesting that dry environments force species with similar ecological traits to coexist in the communities. This has been interpreted as evidence for the effect of environmental filtering (Graham et al. 2009, Graham et al. 2012). Also, Gomez *et al* (2010), demonstrated that at the regional scale, antbird assemblages are most likely structured by the environment. Species that coexist in ecoregions were shown to have similar traits related to physiological tolerances (Gomez et al. 2010).

The Magdalena Valley has a complex history of climate fluctuations and presumed community composition. During the Pleistocene, the valley was a continuous dry forest that connected the south end with the Caribbean lowlands in Colombia (Haffer 1967). Interestingly, dry forest birds that became isolated from the Caribbean lowlands have successfully immigrated or persisted in the edges of the wet forests (Gomez, J.P. *pers. obs*.). Many of the birds that can be considered as dry forest indicators successfully invade the edges and the surroundings of the wet forest end of the valley (e.g. *Conirostrum leucogenys, Euphonia concinna*). The reason why they are not reflected in the analysis is because the point counts are located inside the forest. These birds are mainly found in scattered trees in pastures, live fences and small remnants of forest (~<1 hectare), where typical wet forest birds are not found. This means that they can tolerate the conditions of the wet forests but their establishment inside the forest is limited by other mechanisms.

Colombia has more bird species than any other country in the world, and our data suggest that moisture gradients play a crucial role in maintaining as well as generating this diversity. Much of this regional diversity may result from dispersal limitation through geographic isolation as a product of Colombia’s complex geography (Smith et al. 2014). Yet, our results suggest that moisture gradients, which are ubiquitous in Colombia, may contribute greatly to this diversification. Most elevation gradients in the Andes also show strong precipitation gradients. The harsh environments created by the extremes of these gradients likely act as stronger filters than either variable acting independently. If threshold effects such as those we observed in the Magdalena Valley are a general phenomenon, then we might observe similar community changes on even smaller spatial scales along these moisture/temperature gradients, a result that should generate rapid community turnover and very high beta diversity.

## Acknowledgements

We thank Hda. Los Limones, C. Garcia, C. Mendoza, H. Llara, Remanso del Sumapaz, Pizano-Gomez Family, A.M. Jaramillo, Rio Claro, J.L. Toro, Corantioquia, A. Link, O. Laverde and G. Campuzano for support in the study sites. J. Drucker, A. Echeverri, J. Llano-Mejia, M.A. Loaiza, A. Morales-Rozo, E. Ortiz-Acevedo, J.L. Parra, J. Sandoval and E. Yepes for their assistance in the field. J. Ferguson and J.M. Ponciano for stimulating discussions and comments on the manuscript. Funding sources Chapman AMNH, COS, Sigma-Xi and National Geographic-Waitts Grant No. W270-13 to J.P Gomez and NSF grant DEB 213858 to S.K. Robinson.

---

**Table S1.**
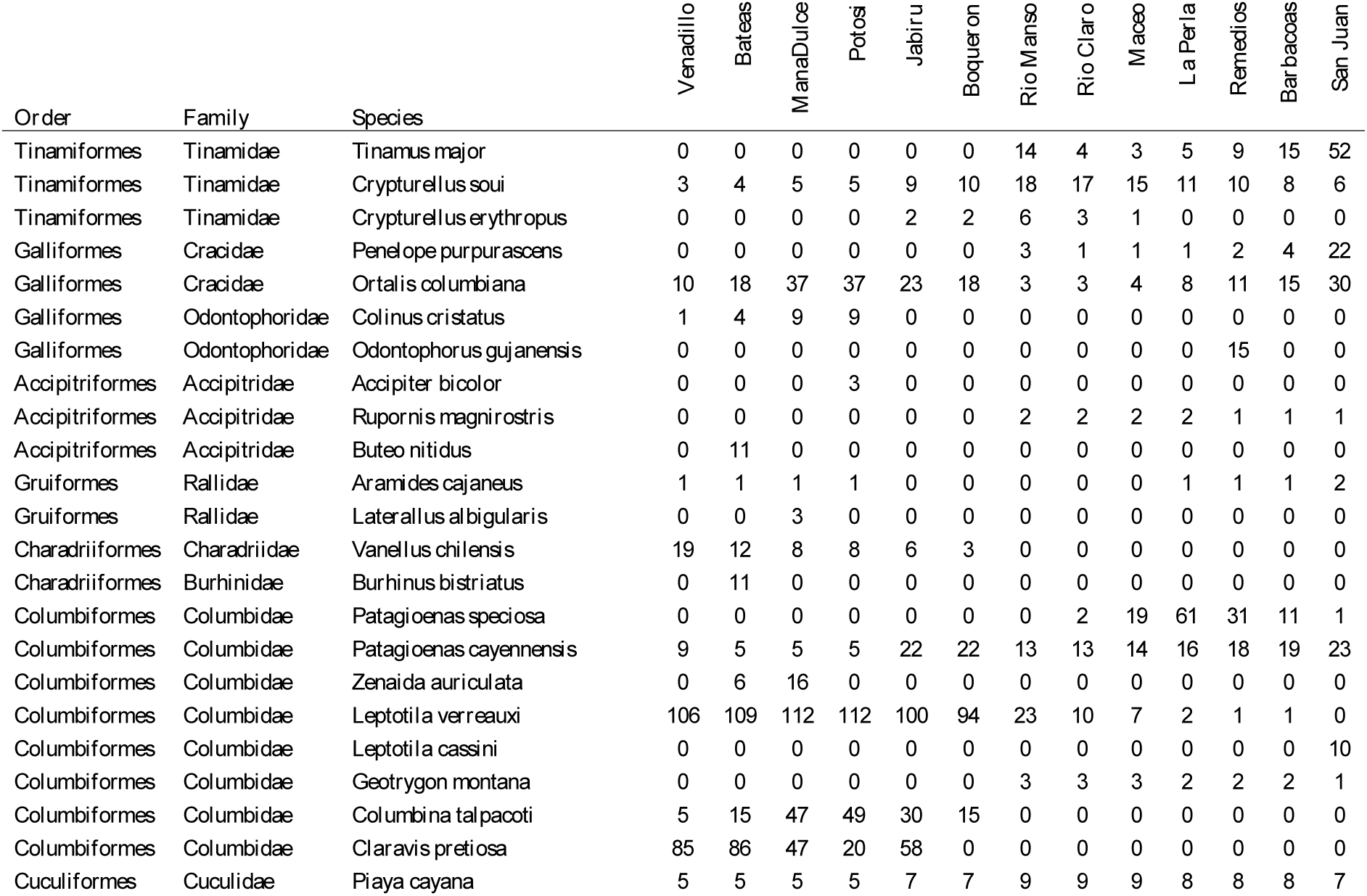
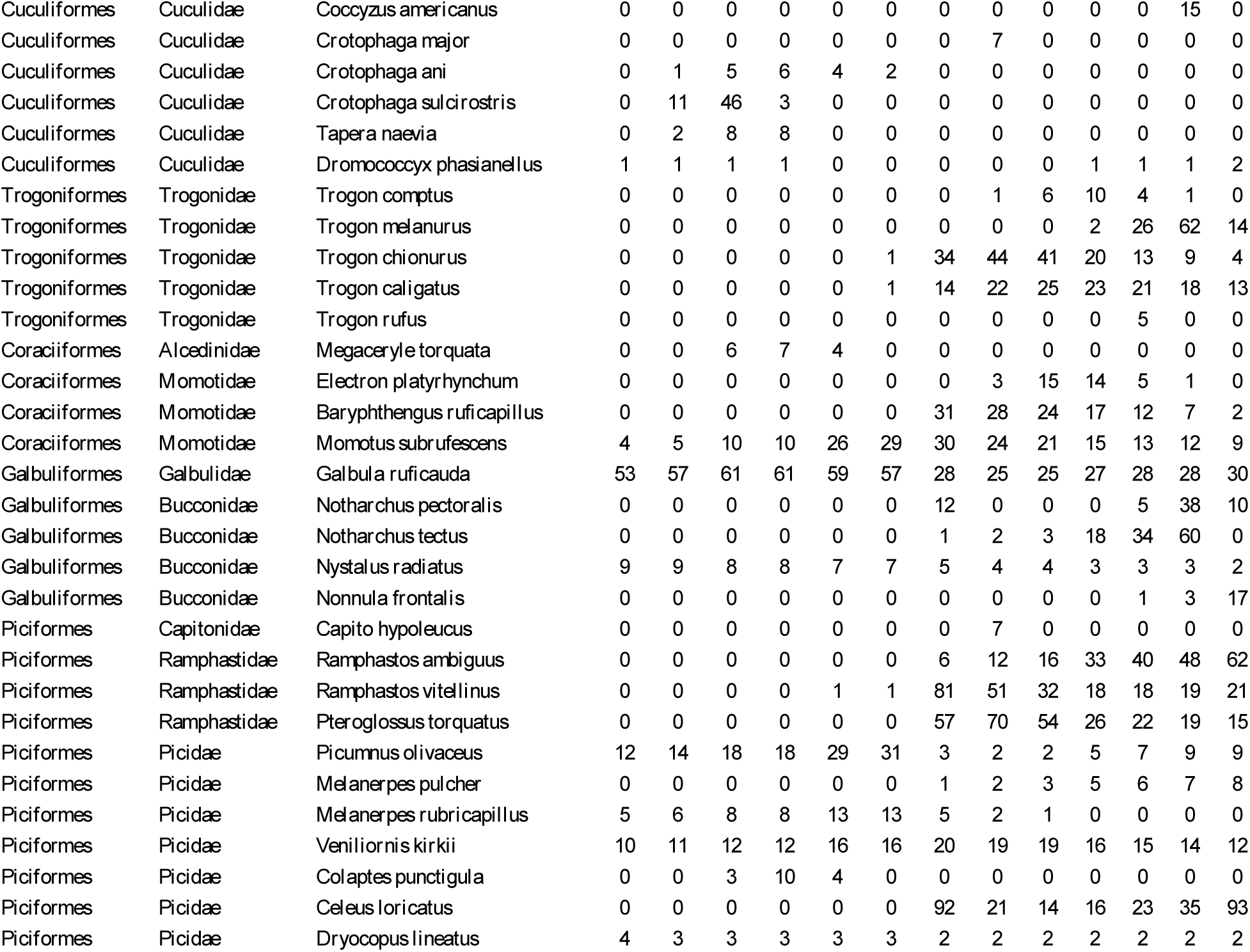
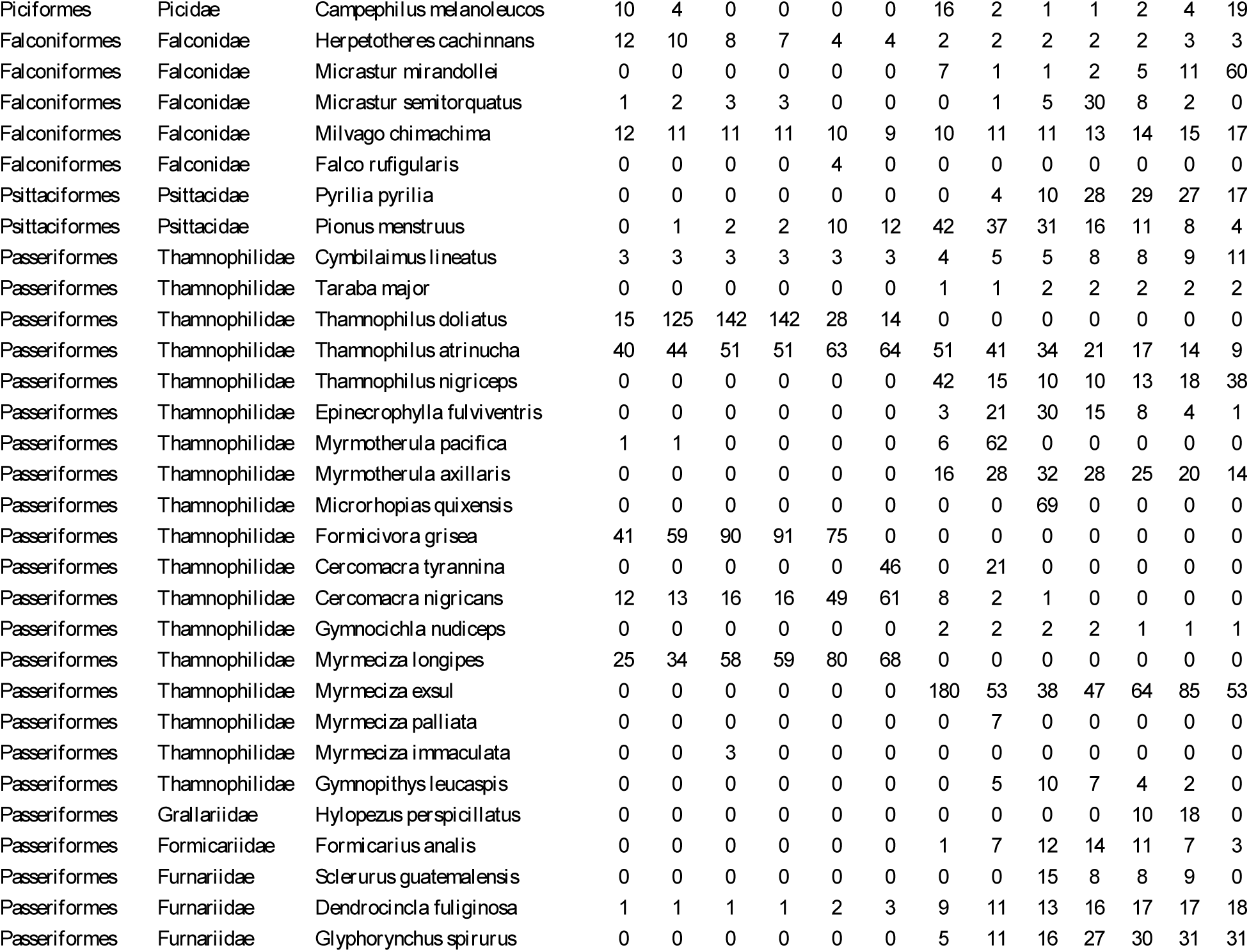
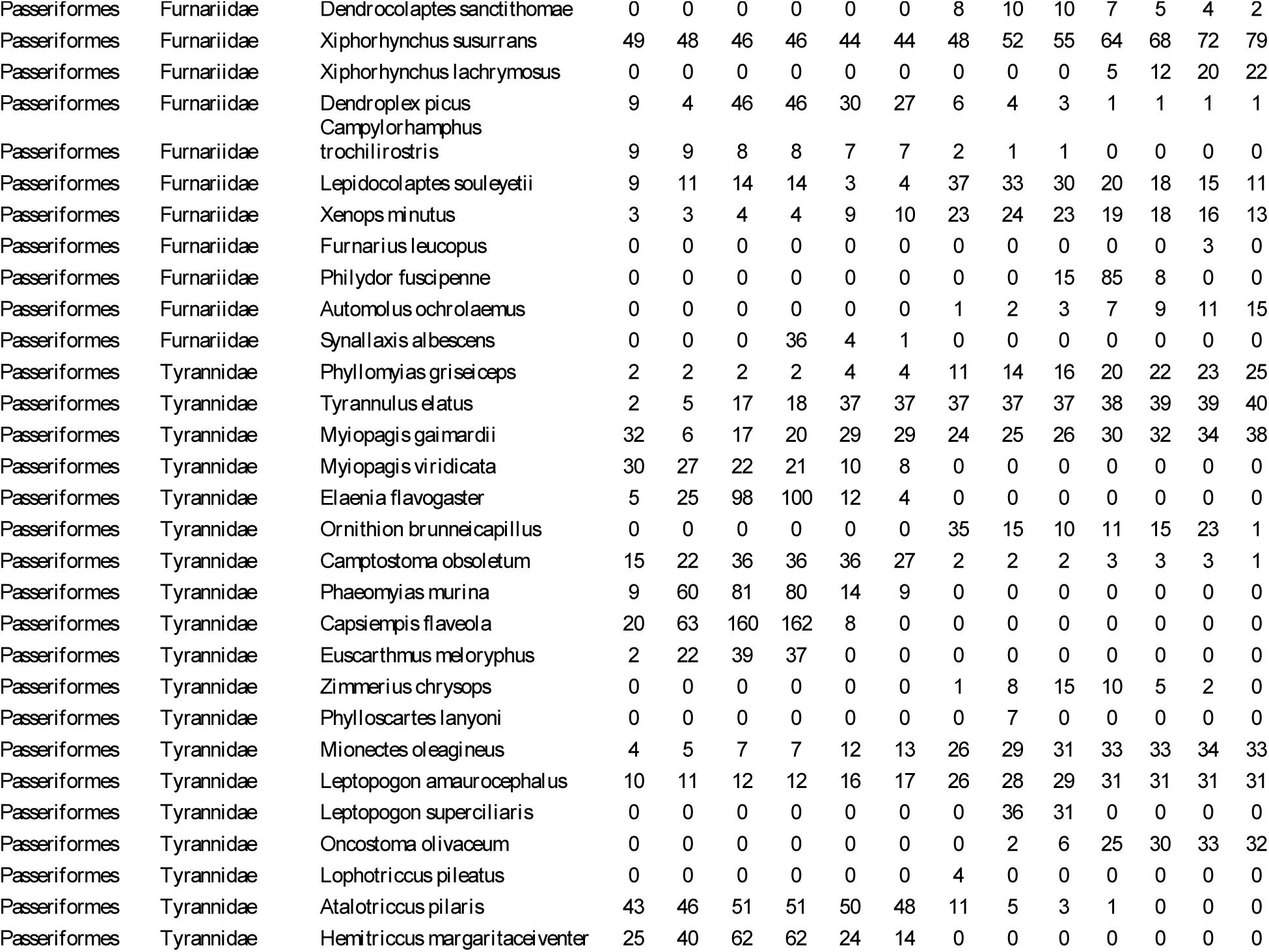
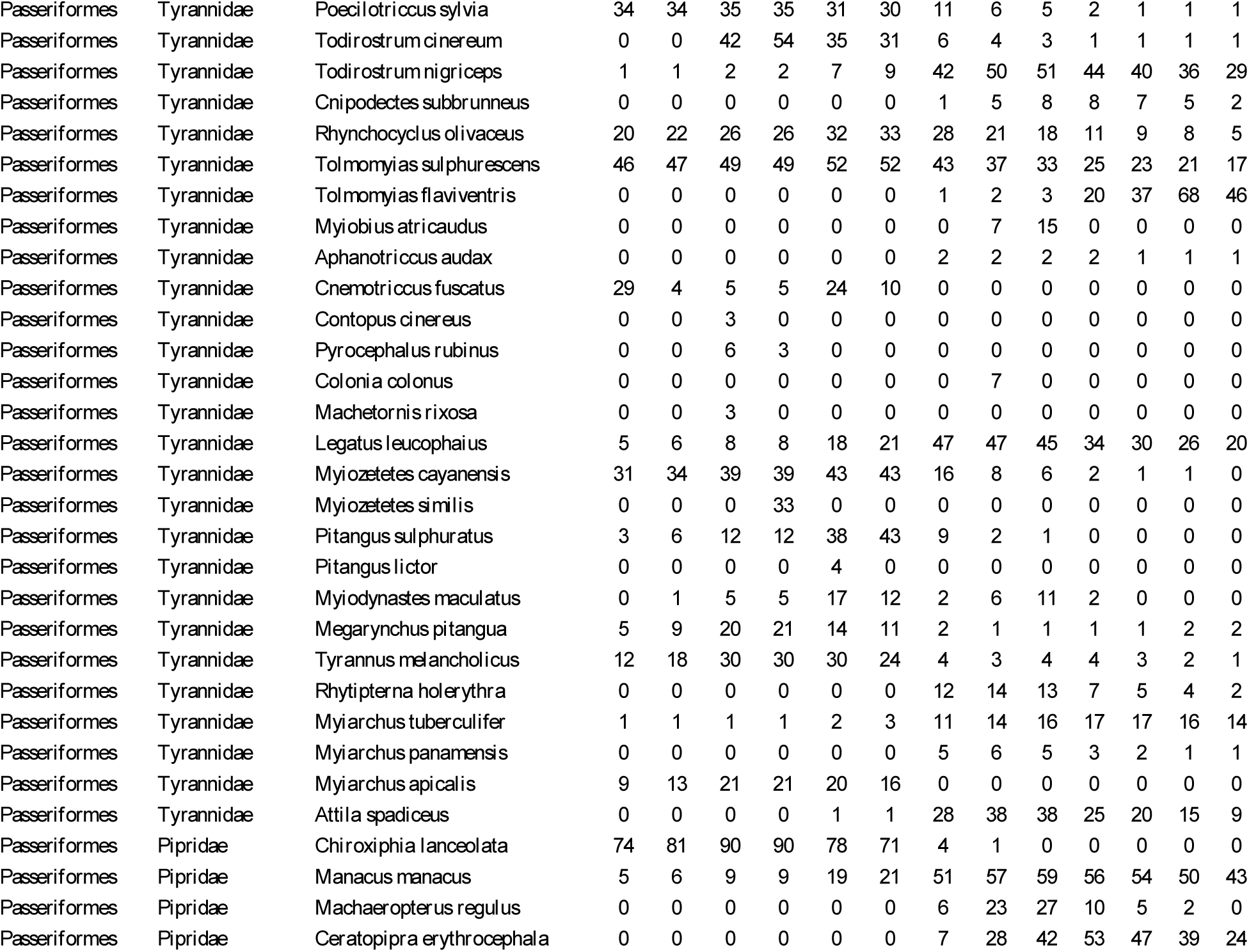
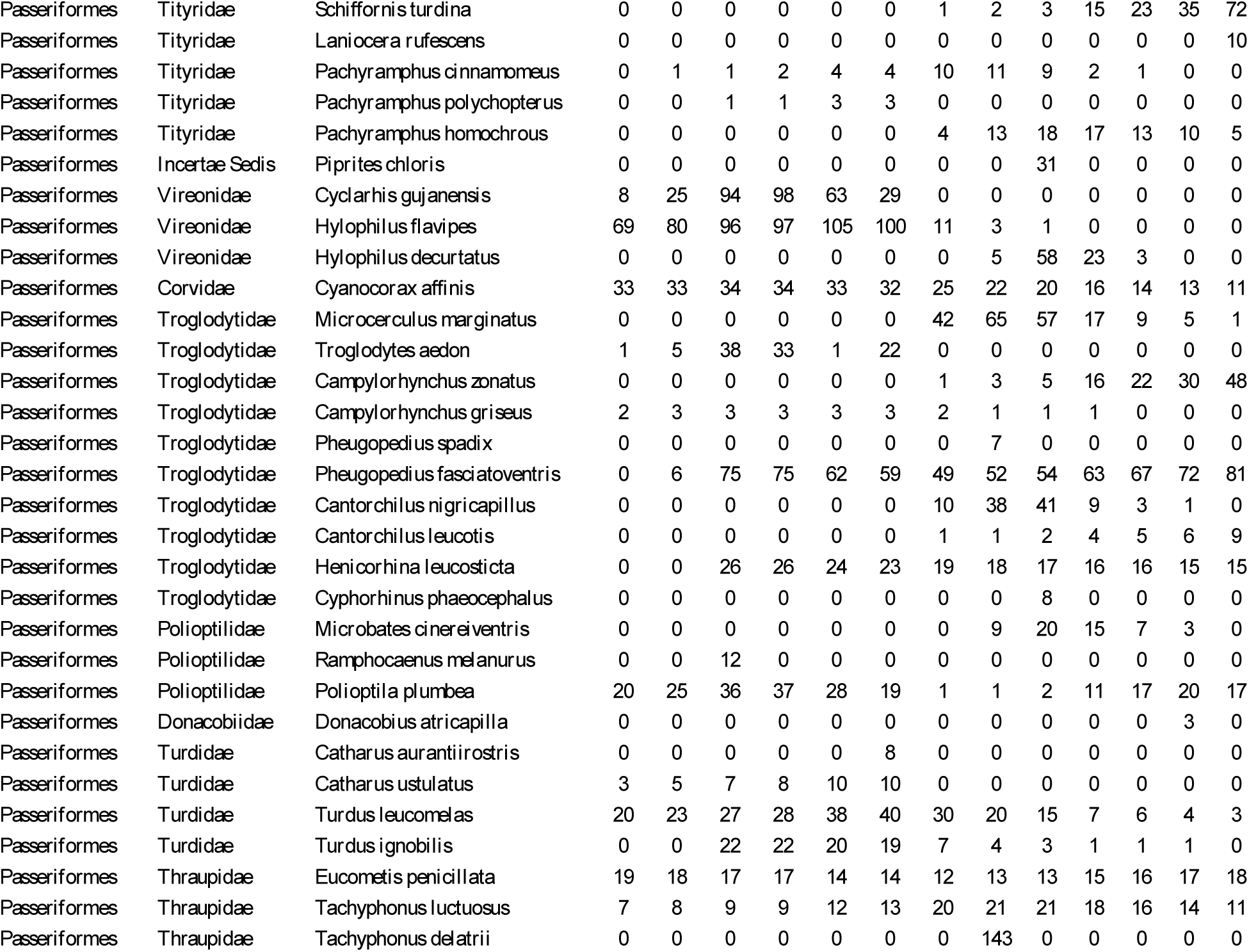
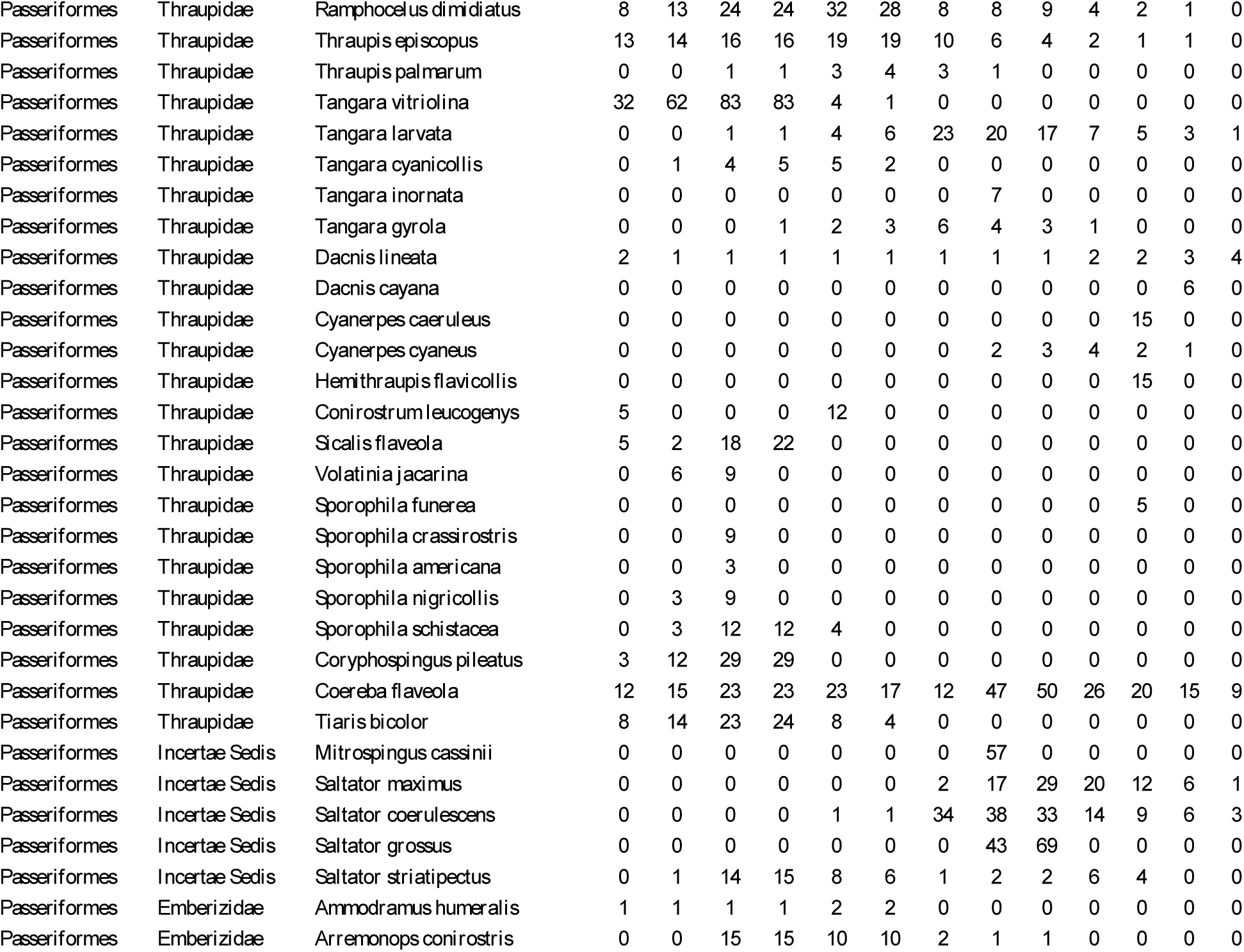
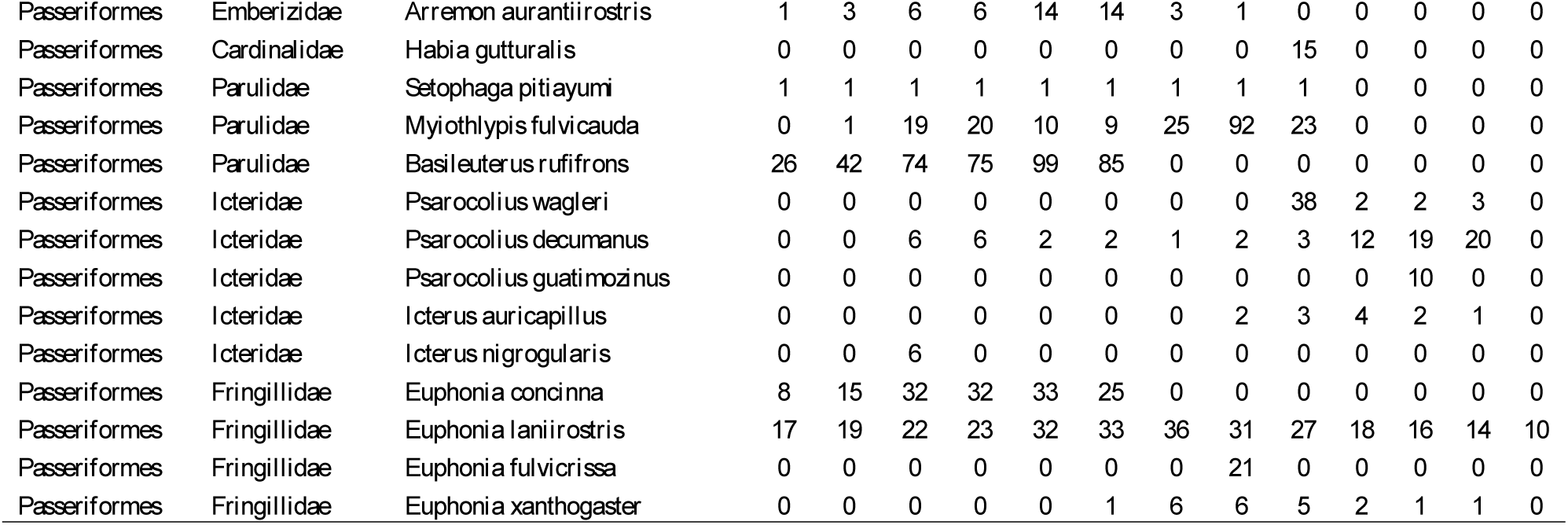
Population density estimates for species in each of the 13 localities. The estimates are the results from the Zero-Inflated Poisson model and extrapolated to 100ha. The population of each species in each local community is given in individuals/100ha

**Table S2.**
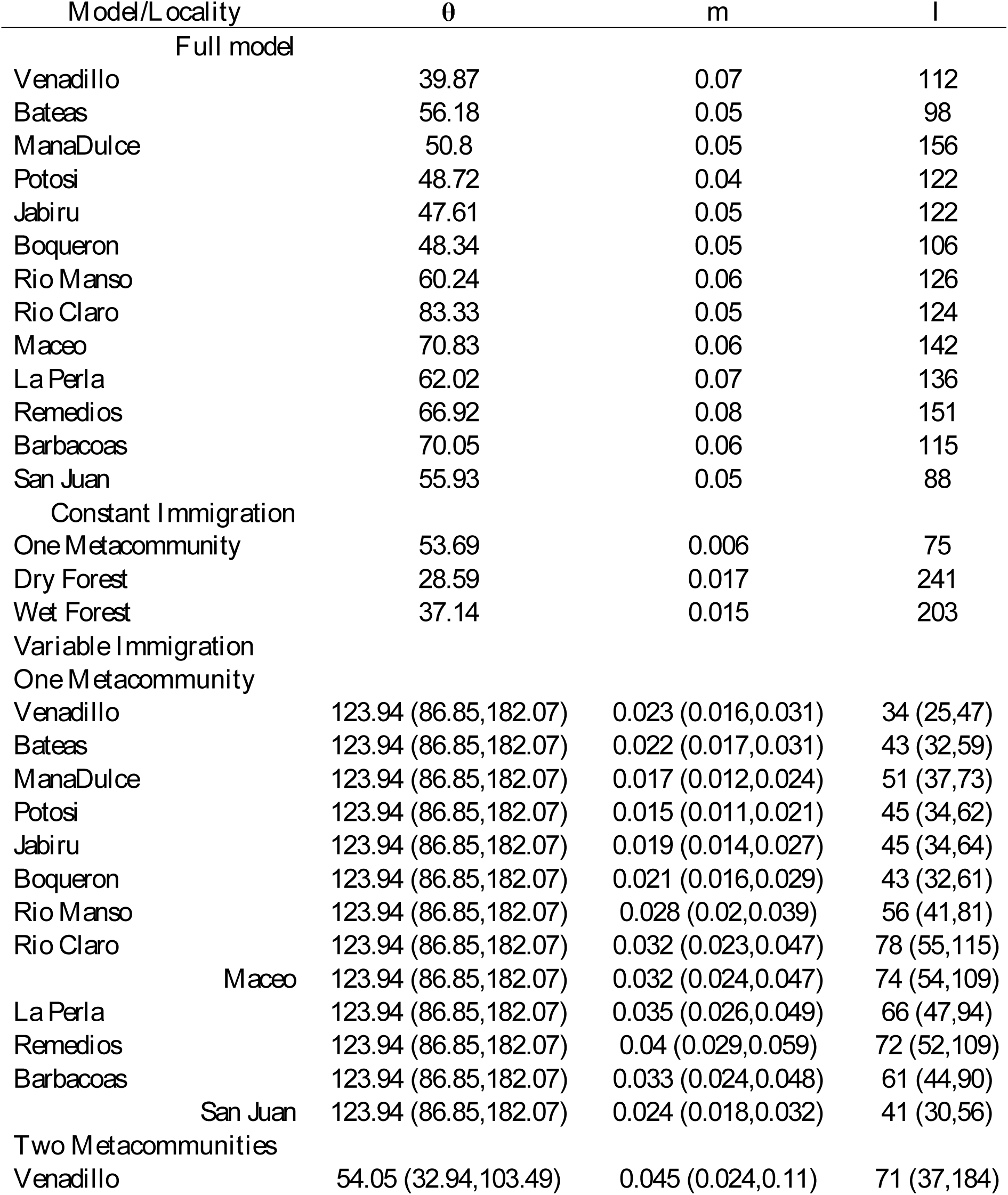
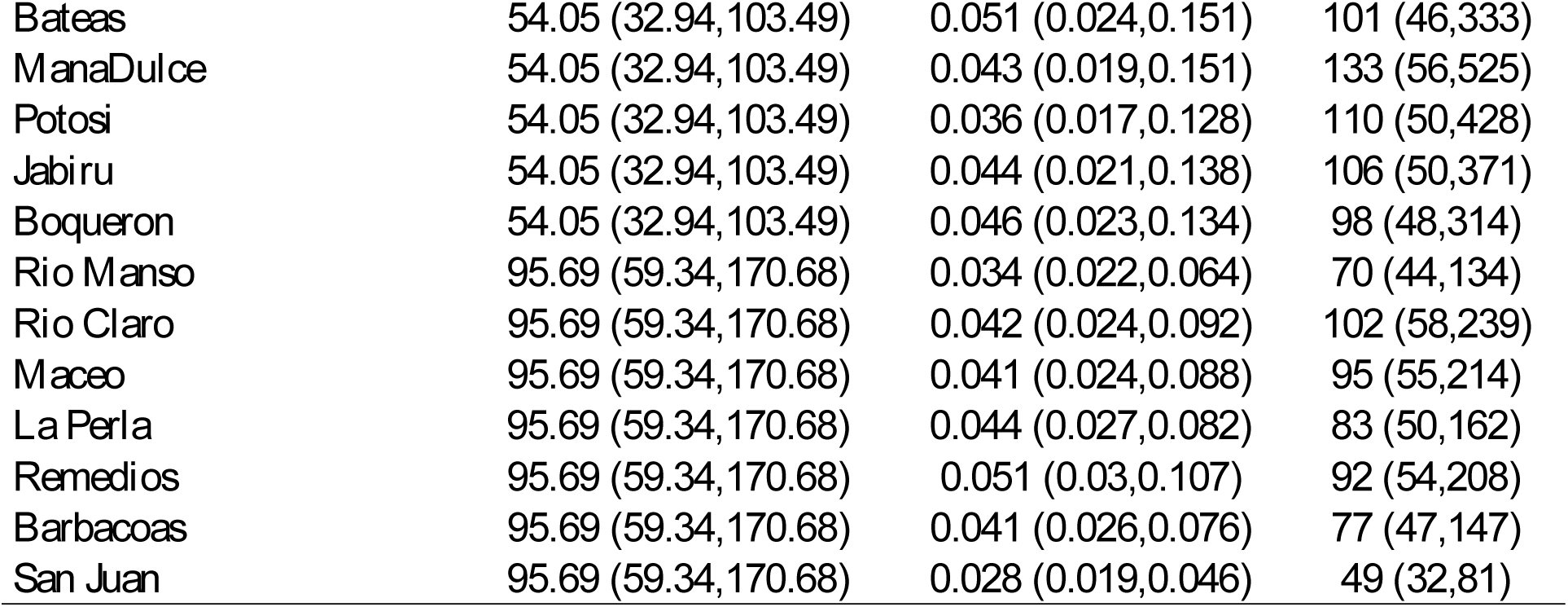
Results of the Maximum likelihood estimation of the parameters for each of the models tested. We present parametric bootstrap confidence intervals only for the Etienne 2009 model. The results from the Neutrality tests are also shown.

**Table S3.** Results of the models of the relationship between Immigration Rate (m), Potential Immigrants (I) and Precipitation and distance from the centroid of the metacommunity. Terms in bold indicate significance at the 0.05 level.

| Model | Intercept | Precipitation | Distance | p-value | R<sup>2</sup> | AIC | ΔAIC |
| --- | --- | --- | --- | --- | --- | --- | --- |
| *One Metacommunity* |  |  |  |  |  |  |  |
| *m* |  |  |  |  |  |  |  |
| Full | 0.005 | **0.009** | 6.92E-06 | >0.01 | 0.68 | -97.40 | 1.87 |
| Precipitation | 0.007 | **0.009** |  | >0.01 | 0.67 | -99.27 | 0.00 |
| Distance | **0.030** |  | -2.97E-05 | 0.37 | 0.07 | -85.75 | 13.51 |
| *I* |  |  |  |  |  |  |  |
| Full | **38.81** | **11.94** | -7.05E-02 | 0.01 | 0.59 | 101.42 | 0.00 |
| Precipitation | **23.68** | **14.79** |  | 0.01 | 0.49 | 102.27 | 0.84 |
| Distance | **69.77** |  | **-1.17E-01** | 0.04 | 0.32 | 106.08 | 4.66 |
| *Two Metacommunities* |  |  |  |  |  |  |  |
| *m* |  |  |  |  |  |  |  |
| Full | **0.05** | 0.00 | -4.40E-05 | 0.43 | 0.15 | -90.07 | 0.94 |
| Precipitation | **0.05** | 0.00 |  | 0.34 | 0.08 | -91.01 | 0.00 |
| Distance | **0.04** |  | -2.73E-05 | 0.57 | 0.03 | -90.29 | 0.72 |
| *I* |  |  |  |  |  |  |  |
| Full | **128.21** | -15.56 | -5.43E-02 | 0.28 | 0.23 | 120.27 | 211.28 |
| Precipitation | **0.05** | 0.00 |  | 0.34 | 0.08 | -91.01 | 0.00 |
| Distance | **0.04** |  | -2.73E-05 | 0.57 | 0.03 | -90.29 | 0.72 |

**Table S4.** Results of the models of the relationship between number of rare species and Precipitation and total number of species. The coefficients are the result of a Poisson regression in the form: # *Rare spp* = *e*^*A*+*β*_1_*x*_1_+*β*_2_*x*_2_^ where A is the intercept of the model, *x*_1_ and *β*_1_ are Precipitation and its estimated coefficient and *x*_2_ and *β*_2_ are Richness and its coefficient. In this case three levels of rarity where considered: less than 2 individuals/Ha, less than 5 individuals/Ha and less than 10 individuals/Ha. Terms in bold indicate significance at the 0.05 level.

| Model | Intercept | Precipitation | Richness | LogLik | AIC | ΔAIC |
| --- | --- | --- | --- | --- | --- | --- |
| **2 ind/Ha** |  |  |  |  |  |  |
| *Richness* | **1.38** |  | **0.02** | -41.00 | 86.00 | 6.56 |
| *Precipitation* | **2.23** | **0.41** |  | -37.80 | 79.60 | 0.16 |
| *Full* | **1.61** | **0.33** | 0.01 | -36.72 | 79.44 | 0.00 |
| **5 ind/Ha** |  |  |  |  |  |  |
| *Richness* | **2.90** |  | **0.006** | -44.32 | 92.66 | 3.40 |
| *Precipitation* | **3.25** | **0.2** |  | -42.62 | 89.25 | 0.00 |
| *Full* | **3.02** | **0.16** | 0.002 | -42.34 | 90.68 | 1.42 |

